# Optimal Performance Objectives in the Highly Conserved Bone Morphogenetic Protein Signaling Pathway

**DOI:** 10.1101/2024.02.01.578451

**Authors:** Razeen Shaikh, Nissa J. Larson, Donny Hanjaya-Putra, Jeremiah Zartman, David M. Umulis, Linlin Li, Gregory T. Reeves

**Affiliations:** Artie McFerrin Department of Chemical Engineering, Texas A&M University, College Station, TX 77843, United States; Weldon School of Biomedical Engineering, Purdue University, West Lafayette, IN 47907, United States; Aerospace and Mechanical Engineering, University of Notre Dame, Notre Dame, IN 46556, United States; Bioengineering Graduate Program, University of Notre Dame, Notre Dame, IN 46556, United States; Chemical and Biomolecular Engineering, University of Notre Dame, Notre Dame, IN 46556, United States; Faculty of Genetics and Genomics, Texas A&M University, College Station, TX 77843, United States

## Abstract

Throughout development, complex networks of cell signaling pathways drive cellular decision-making across different tissues and contexts. The transforming growth factor β (TGF-β) pathways, including the BMP/Smad pathway, play crucial roles in these cellular responses. However, as the Smad pathway is used reiteratively throughout the life cycle of all animals, its systems-level behavior varies from one context to another, despite the pathway connectivity remaining nearly constant. For instance, some cellular systems require a rapid response, while others require high noise filtering. In this paper, we examine how the BMP-Smad pathway balances trade-offs among three such systems-level behaviors, or “Performance Objectives (POs)”: response speed, noise amplification, and the sensitivity of pathway output to receptor input. Using a Smad pathway model fit to human cell data, we show that varying non-conserved parameters (NCPs) such as protein concentrations, the Smad pathway can be tuned to emphasize any of the three POs and that the concentration of nuclear phosphatase has the greatest effect on tuning the POs. However, due to competition among the POs, the pathway cannot simultaneously optimize all three, but at best must balance trade-offs among the POs. We applied the multi-objective optimization concept of the Pareto Front, a widely used concept in economics to identify optimal trade-offs among various requirements. We show that the BMP pathway efficiently balances competing POs across species and is largely Pareto optimal. Our findings reveal that varying the concentration of NCPs allows the Smad signaling pathway to generate a diverse range of POs. This insight identifies how signaling pathways can be optimally tuned for each context.

## Introduction

Throughout the process of development, an array of essential communication pathways exert influence over a wide spectrum of cellular destiny determinations and various processes in different tissues and situations, including the Bone Morphogenetic Protein (BMP) pathway, a member of the Transforming Growth Factor β (TGF-β) superfamily of signaling pathways. The BMP pathway regulates a wide variety of cellular responses, including apoptosis, differentiation, homeostasis, stem cell maintenance, and regeneration in animals from flies to humans [1–7]. The pathway is activated by ligands of the BMP family, which bind to cognate Type I serine-threonine kinase receptors, promoting the recruitment of the Type II receptors (Fig. 1A). The Type I/Type II receptor complex phosphorylates Smad1/5/8, which then dimerizes and forms a complex with the Co-Smad (Smad4), forming the signaling complex (PSmad1)_2_/Smad4. This signaling complex (SC) enters the nucleus and activates downstream gene expression [1, 8]. The first Smads were discovered in *Drosophila*: Mothers against Dpp (Mad; homolog of Smad1/5/8) and Medea (Med; homolog of Smad4) [9].

**Figure 1:**
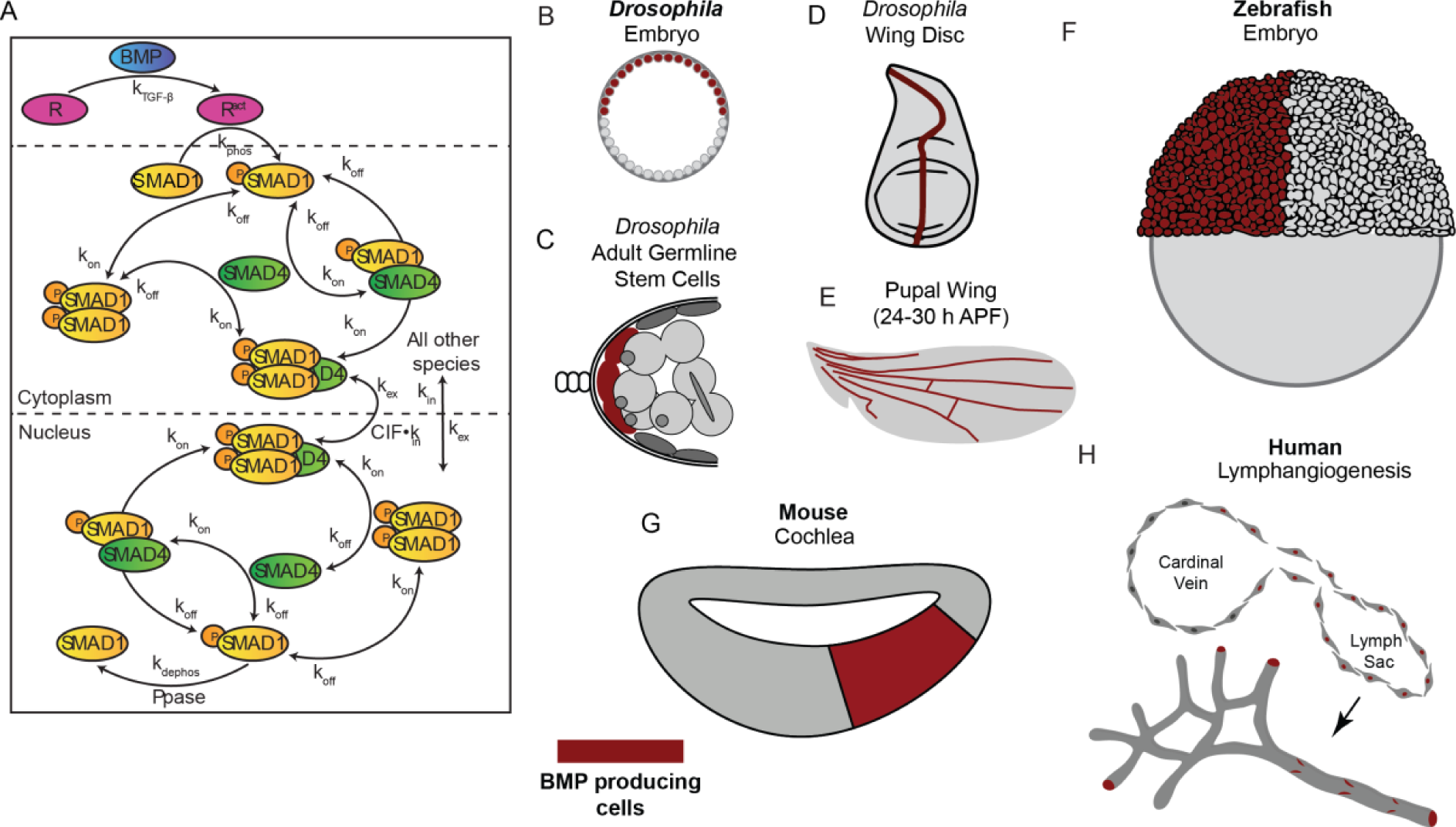
Smad signaling is a conserved pathway. (A) Reaction network diagram of the Smad pathway. TGF-β binds and activates the receptors, thus phosphorylating Smad1. pSmad1 binds to Smad4 and forms dimers which then form trimers. **(B- H)** Illustrative figures of model systems in which BMP signaling pathway patterns tissues with BMP producing cells marked in maroon, **(B)** *Drosophila* Embryo, **(C)** *Droso*phila adult germline stem cells, **(D)** *Drosophila* Wing Disc, **(E)** *Drosophila* Pupal Wing (24-30 h APF), **(F)** Zebrafish Embryo, **(G)** Mouse Cochlea, and **(H)** Human Lymphangiogenesis

Despite its high degree of conservation across different species, the BMP/Smad pathway exhibits remarkable diversity in its responses to BMPs in different contexts. For example, the response speed of the Smad pathway varies widely across common model systems, including several well-studied stages of *Drosophila* development, the zebrafish embryo, the mammalian cochlea, and human induced pluripotent stem cells (hiPSCs) (Fig. 1B-H).

For example, BMP signaling patterns the dorsal half of the embryo in the 2-3 h old *Drosophila* embryo, [10, 11] to establish a narrow profile in the dorsal-most of the embryo in the first 30 min [12–15]. Thus, the *Drosophila* Smad network rapidly responds in the embryo. Later in development, the same Smad pathway acts on the order of hours in the specification of wing veins [16]. In the case of vertebrate development, the zebrafish embryo BMP gradient is formed on a rapid time scale (30 min) [17, 18]. Whereas a similar pathway responds to the different systems in a varied time scale. In contrast, BMP signaling in lymphatic commitment in humans is on the order of hours, so that noise filtering is a more dominant performing objective than response speed [19–22]. As demonstrated in each of these model systems, the time scale of BMP pathway activation and response, and noise control response to the input signal must be tightly regulated.

These differences in response speed highlight the fact that the BMP-Smad pathway exhibits a diversity of behavior depending on the biological context. It is likely that the response speed is in a trade-off relationship with other systems-level behaviors, or “Performance Objectives (POs)” [23, 24]. For example, communication systems can sacrifice response speed to reduce noise through time-averaging [25, 26]. To simultaneously improve noise filtering and response speed, signaling pathways could operate in the regime of high levels of activated receptor inputs. However, in this regime, linear sensitivity to input levels is sacrificed, as the network would become saturated [25, 26]. Indeed, the utopian ideal — a signaling system with an instantaneous response, zero noise, and a perfectly linear, high-fidelity response — is not biologically or physically possible, and compromises must be made to balance the trade-offs among the POs. Certain cellular systems might demand swift responses, like *Drosophila* and zebrafish embryos, whereas others may necessitate extensive noise filtration, like human stem cells. This diversity cannot derive from rewiring the pathway, or from altering protein biochemical function, as these aspects of the BMP pathway are highly conserved across the animal kingdom. Instead, we hypothesize the tunability is achieved through differential concentrations of pathway components.

In this study, we used a Smad pathway model, calibrated to time course data from human cells [27], to analyze how the Smad pathway balances trade-offs among the three POs through variation of non-conserved parameters (NCPs): concentrations of phosphatase and Smad proteins, and the nuclear import rate of activated Smad protein complex. We found that the NCPs allow the Smad signaling pathway to generate a diverse range of POs, and that phosphatase concentration has the largest effect on shifting the balance of POs. To systematically determine which combinations of POs are optimal, we utilized the concept of the Pareto Front, which has been widely employed in economics to identify a collection of designs that offer optimal trade-offs among various requirements [28, 29]. In the field of biology, the Pareto Front approach allows for a comprehensive analysis of the diverse factors influencing biological systems and aids in the identification of solutions that achieve the best compromise between competing objectives. We found that, for most combinations of NCP values, the Smad pathway is Pareto optimal. We analyzed which systems fall on the Pareto Front and predicted the relationship between values of NCPs and which POs a given system would emphasize. We conclude that the Smad pathway is highly versatile to achieve a variety of trade-offs among the POs, depending on the needs of each system in their biological context.

## Results

### The non-conserved parameters determine the performance objectives of the system

Our BMP/Smad pathway model was adapted from a similar model constructed for the Smad pathway downstream of the TGF-β pathway [27]. We used a core set of parameters that were fit to the TGF-β/Smad data from [27]. With the core parameters in place, we performed simulations of the BMP/Smad pathway (hereafter referred to simply as the Smad pathway) with the pathway initially at rest (active receptor level, R_act_ = 0). At time t = 0, we increased the active receptor level to its new value, R_act_, which resulted in an increase in the nuclear concentration of the signaling complex (SC), (PSmad1)_2_/Smad4. Based on this simulation, we computed the rise time (t_rise_), which serves as an indicator of the system’s response speed (PO1). The t_rise_ is defined as the duration it takes for the response variable to reach and consistently stay within a 5% window around the final steady state (Fig. 2A).

**Figure 2:**
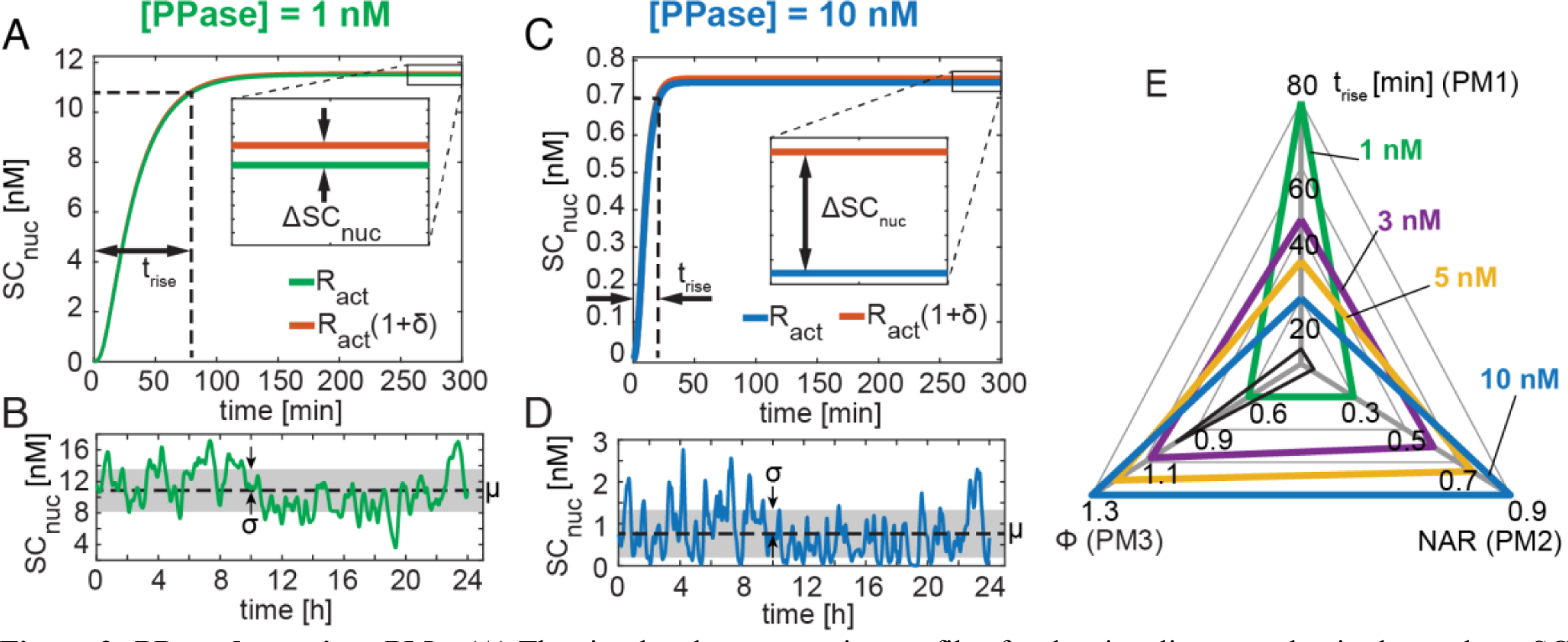
PPase determines PMs: (A) The simulated concentration profiles for the signaling complex in the nucleus, SC_nuc_, at a constant activated receptor level and PPase concentration of 1 nM, has a t_rise_ of 80 min and a small effect on the steady- state concentration when the R_act_ level is perturbed. **(inset)** The differential change in steady-state concentration with respect to change in R_act_ is 0.6. **(B)** In response to the noise input (S5A) the resulting dynamic profile for SC_nuc_ has a small coefficient of variation at PPase concentration of 1 nM (black dashed line indicates mean and grey rectangle indicates standard deviation). **(C)** The simulated concentration profile for SC_nuc_ at a constant activated receptor level and PPase concentration of 10 nM has a t_rise_ of 20 min and a relatively (relative to 1 nM) larger change in steady-state concentration when perturbed. **(inset)** The differential change in steady-state concentration with respect to change in R_act_ is 1.3. **(D)** In response to the noise input (S5A) the resulting dynamic profile for SC_nuc_ has a large coefficient of variation at PPase concentration of 10 nM. **(E)** The Performance Metrics (PMs) of the Smad signaling pathway depend on the PPase concentration. With an increase in PPase concentration from 1 nM to 10, nM t_rise_ (PM 1) decreased, NAR (PM 2) increased, and sensitivity coefficient (φ) (PM 3) increased. The PPase concentrations for which the PMs are calculated as 1 nM, 3 nM, 5 nM, and 10 nM. The “utopian ideal”, in which all three PMs are optimized, is shown in black

To evaluate the noise amplification properties of the Smad pathway, we extended the simulation, beginning at the established steady state, but with a time-varying noise R_act_ (Fig. S5A). The variations in R_act_ mimic stochastic simulations of receptor activation [30]. The simulation was continued over a simulation time of 24 hours. During that time, the response variable fluctuated about a mean of μ with a standard deviation of σ (Fig. 2B). Using these simulated fluctuations in the response variable, we calculated the second metric, NAR [31], which is defined as the ratio of the coefficient of variation 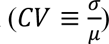 of the response variable tohat of the input variable (R_act_). The final metric, the steady state sensitivity coefficient (ϕ), is a measure of the linear fidelity of the response variable with respect to the input variable. To calculate this metric, we ran a parallel simulation by perturbing R_act_ by 1%, resulting in a slightly different steady state for the response variable (inset of Fig. 2A). The sensitivity coefficient is defined as the ratio of the fractional change in steady state output to that of the input.

Each of these three performance metrics (PMs) was calculated for a given set of values of the NCPs, and in general, should change when the values of the NCPs change. Fig. 2E shows how the PMs change while perturbing phosphatase (PPase). We found that increasing the PPase concentration from 1 nM to 10 nM, keeping all other core parameters and NCPs unchanged, resulted in a decrease in the rise time (Fig. 2A, C), an increase in NAR (Fig 2B, D), and an increase in the sensitivity coefficient (inset of Fig. 2A, C). These trends were also maintained for intermediate values of the phosphatase concentration (Fig. 2E). By comparison, the utopian ideal has t_rise_ = 0, NAR = 0, and ϕ = 1 (black triangle in Fig. 2E), which is not biologically or physically possible. In summary, our results indicate that the PPase concentration affects the performance measures (PMs). Therefore, by regulating PPase concentration, the Smad signaling pathway can be fine-tuned to achieve specific desired outcomes.

### The effect of non-conserved parameters on the performance objectives

While the Smad pathway topology and component protein sequence is highly conserved across the animal kingdom, we consider four parameters in the model to be “non conserved parameters (NCPs)”. The first three are the concentrations of the PPase (see above), Smad1, and Smad4. The fourth NCP in the model is the complex import factor (CIF), which is defined as the nuclear import rate constant of the SC normalized to the nuclear import rate of the other Smad-containing complexes (Schmierer et al., 2008).

To determine how each of these NCPs affects the three PMs, we performed a parameter screen and calculated the three PMs for 10^4^ sets of randomly generated sets of NCPs (see Methods) spanning several orders of magnitude. This extensive analysis allowed us to generate a three-dimensional manifold of all feasible PMs in “PM-space” (Fig. 3A), which revealed that no parameter set resulted in optimal values of all three PMs simultaneously (designated as the utopian point, UP). The projections of the three-dimensional manifold indicate that the PM-space is constrained by the model formulation (Fig. 3B-D). Notably, trade-offs were observed between competing objectives such as response speed vs. noise amplification (Fig. 3B), and noise amplification vs. sensitivity coefficient (Fig. 3C).

**Figure 3:**
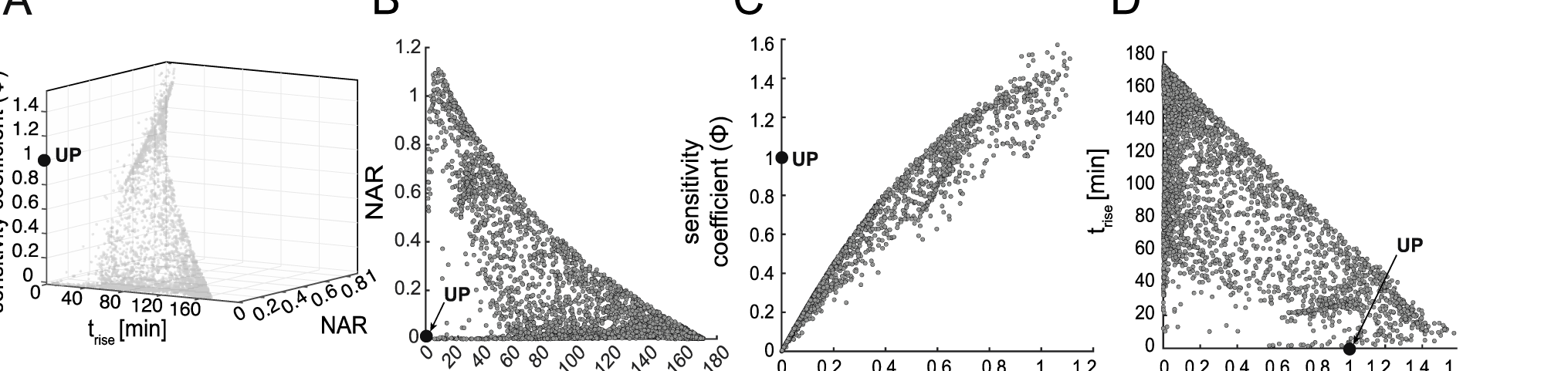
The results of the parameter screen in PM-space. (A) 3D surface representation of the results of the front of the parameter screen in PM-space. Trade-offs are evident as no parameter set results in PMs that approach the utopian point (UP). (B) The rise time (t_rise_)-NAR projection of all PMs generated by sampling NCPs. (C) The NAR-sensitivity coefficient (φ) projections of all PMs generated by sampling NCPs. (D) The sensitivity coefficient (φ) – rise time (t_rise_) projections of all PMs generated by sampling NCPs

To visualize the effect of each NCP on determining the PMs of the signaling network, we sorted the points from the parameter screen into bins based on the value of each NCP (Fig. S6). However, given the high density of points, we replaced each cluster of points by its centroid (see Methods). We found that PPase concentration has the largest effect on the balance of PMs, while CIF has the smallest (Fig. 4A, B). Smad1, Smad4, and their ratio each had a non-monotonic effect on the balance of PMs (Fig. 4C-E).

**Figure 4:**
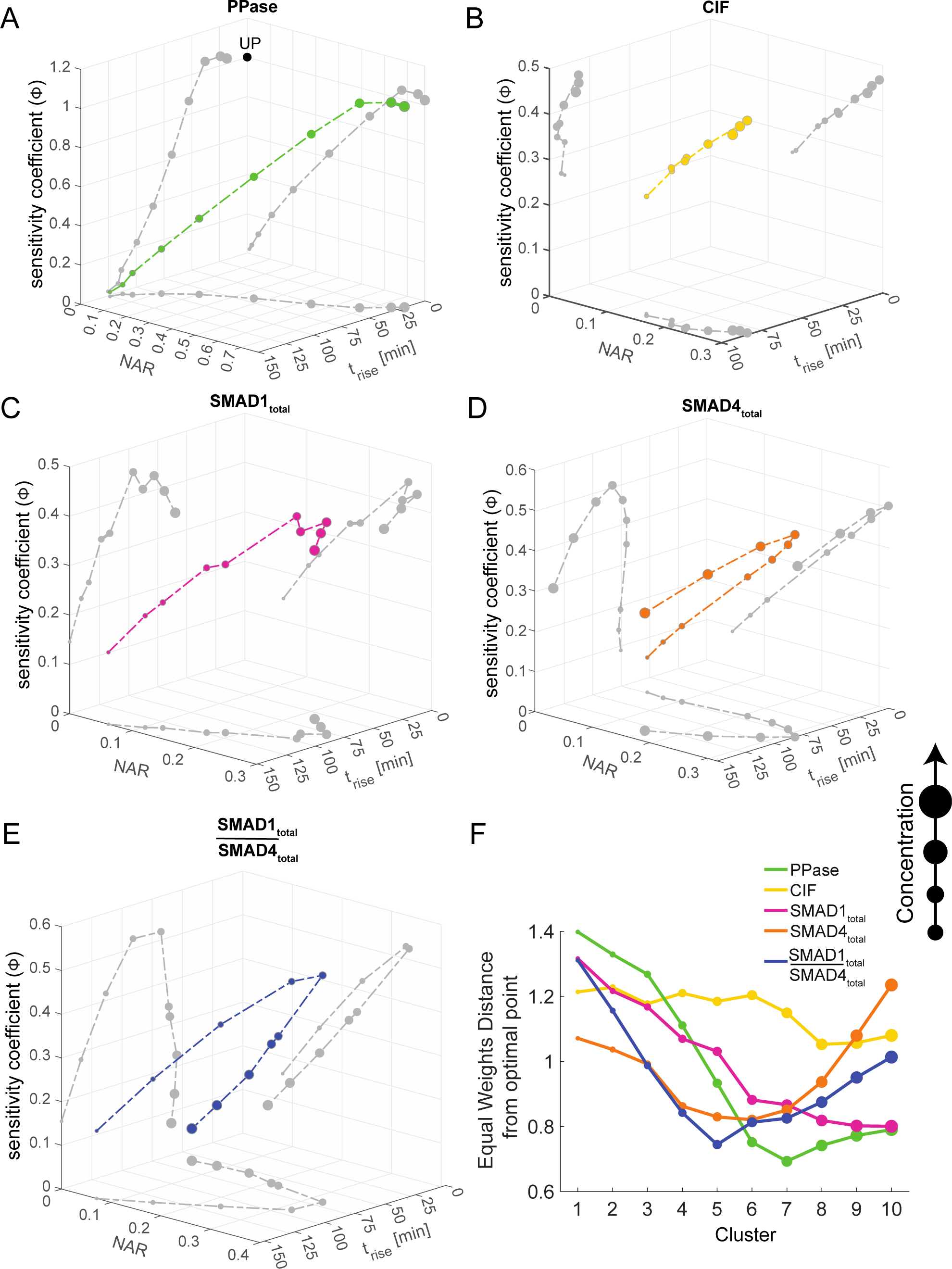
NCPs determine the position of the system in three-dimensional PM-space. (A-E) The centroid of ten point- cloud clusters into which the PMs were sorted based on NCP concentration. The projections of the curve formed by connecting the centroids is as shown on the trise-NAR, trise-sensitivity coefficient (φ), NAR-sensitivity coefficient (φ) planes (shown in grey). Increasing circle diameter denotes increasing concentration. **(A)** An increase in the PPase concentration shifts the PMs of the system closer to the utopian point “UP”. “UP” is the “ideal” optimal performance point with co-ordinates [t_rise_, NAR, φ] = [0, 0, 1] (shown as a black filled circle). **(B)** Increasing CIF concentration has a small effect in determining the position of the centroid of the point cloud clusters. **(C)** Increasing total Smad1 concentration continuously advances the centroid of the point cloud clusters towards the optimal point. After a certain level, any further increase in Smad1 has a small effect in repositioning the PMs. **(D)** The centroid of point cloud clusters forms a “U” shape such that increasing total Smad4 moves the system closer to the optimal point and then after a certain point any further increase in Smad4 moves the system away from the optimal point. **(E)** The Smad1_total_/Smad4_total_ has a mirror “U” trend with lower Smad1_total_/Smad4_total_ values being further away from the optimal than higher Smad1_total_/Smad4_total_. **(F)** The EWD_distance_ of the point cloud cluster from the optimal point for all four NCPs and Smad1_total_/Smad4_total_.

To determine the combination of NCPs for which the system is “optimal,” we calculated the Euclidean distance of each point using equal weights, hereafter referred to as the Equal Weights Distance (EWD) in the curves plotted in Fig. 4A-E to the Utopian Point (UP, Fig. 4A). The utopian points are defined as the “ideal” optimal performance point with minimum rise time (t_rise_ = 0) and noise ratio (NAR= 0), and optimal sensitivity coefficient (φ = 1). While sensitivity coefficient and NAR are order one, unitless ratios , the rise time not unitless and can reach values of a few hundred minutes. To account for this difference in scales, we scaled the rise time by the maximum calculated value over all ten clusters, then arbitrarily weighted the three PMs equally (see Methods). We found there is an optimal level of PPase that minimizes the distance to the utopian point (Fig. 4F). The centroids of CIF clusters are largely equidistant from the utopian point, except at higher concentrations (clusters 8-10) (Fig. 4F). The Smad1 clusters advance towards the utopian point with increasing Smad1 concentration but stagnate such that any further increase in concentration does not significantly affect the position of the clusters (Fig. 4F). In accord with the “U” shape formed by the curves for Smad4 and the Smad1/Smad4 ratio, there are clear optimal values for these NCPs that minimize the distance to the utopian point (Fig. 4F).

### The optimality of performance objectives at different receptor levels

Up to this point, all the simulations described previously were done at a single value of R_act_ (Fig S5A). However, R_act_ levels are expected to vary across species, systems, and space. We generated a total of 60 receptor inputs of varying R_act_ (see Methods) and calculated the PMs at the 10^4^ previously generated NCPs sets. For PPase, at higher R_act_ levels the centroid of point clusters advances closer to the utopian point and stagnates (Fig. 5A). The centroid of point clusters of CIF have the same nature at all R_act_, but the distance from the utopian point progressively increases on increasing R_act_ (Fig. 5B). For all R_act_, increasing total Smad1 shifts the system closer to the utopian point (Fig. 5C) whereas increasing total Smad4 shift the system further away from the utopian point (Fig. 5D). To balance the competing nature of Smad1 and Smad4, the ratio of Smad1_total_/Smad4_total_ must be optimally controlled (Fig. 5E). (Fig. 5 summarizes “R_max_” levels, for “R_min_” see Supplementary Text Section 1).

**Figure 5:**
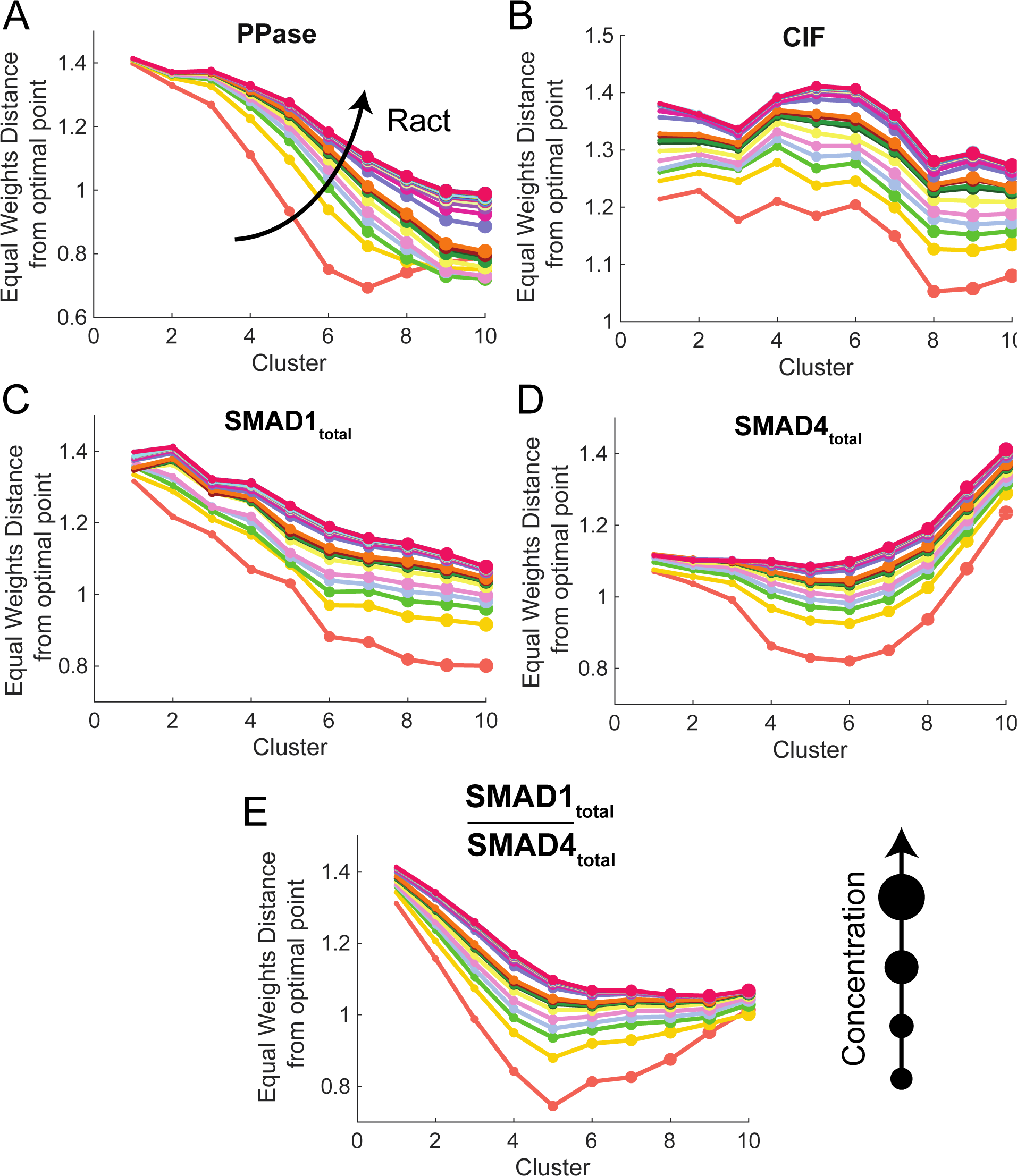
Equal Weights Distance (EWD) distance of point cloud clusters. (A-E) The EWD_distance_ from the optimal point of the point cloud clusters at 30 Rmax (see Methods) inputs. The size of the circle represents increasing NCP concentrations. **(A)** The EWD_distance_ from the optimal point of the point cloud clusters of PPase at 30 R_max_ inputs. **(B)** The EWD_distance_ from the optimal point of the point cloud clusters of CIF at 30 R_max_ inputs. **(C)** The EWD_distance_ from the optimal point of the point cloud clusters of total Smad1 at 30 R_max_ inputs. **(D)** The EWD_distance_ from the optimal point of the point cloud clusters of total Smad4 at 30 R_max_ inputs. **(E)** The EWD_distance_ from the optimal point of the point cloud clusters of total Smad4 at 30 R_max_ inputs.

### The Pareto optimality of the performance objectives

Minimizing the distance of the curves in Fig. 4A-E to the optimal point was based on a EWD_distance_ metric in which the values of t_rise_ (normalized), NAR, and the sensitivity coefficient were given equal numerical weight. However, as the metrics for the three PMs are not related to each other numerically, there is no inherent justification for equal weighting. To address this concern, we have taken a multi-objective optimization (MOO) approach to determine which points in PM-space can be considered optimal. Unlike standard, single-objective optimization, MOO attempts to satisfy multiple, conflicting objectives, and therefore, MOO is the proper method to apply to balance the PMs [32].

In general, in a MOO approach, one identifies the subset of model results in which improvements to one objective cannot be made without compromising the others. This subset is known as the “Pareto Front,” and each of the points on the Pareto Front is considered optimal, from a Pareto standpoint. In practice, points on the Pareto Front are identified as those for which no other point is better in all three objectives. Applying this method to our screen, we found that roughly 70% of results from our screen resided on the Pareto front (Fig. 6). Even some points that perform very poorly on one PM may still considered Pareto optimal because the trade-offs are such that the other PMs are close to their respective optimal values. Therefore, the Smad pathway is tunable to the needs of each system simply by altering the values of each system’s NCPs.

**Figure 6:**
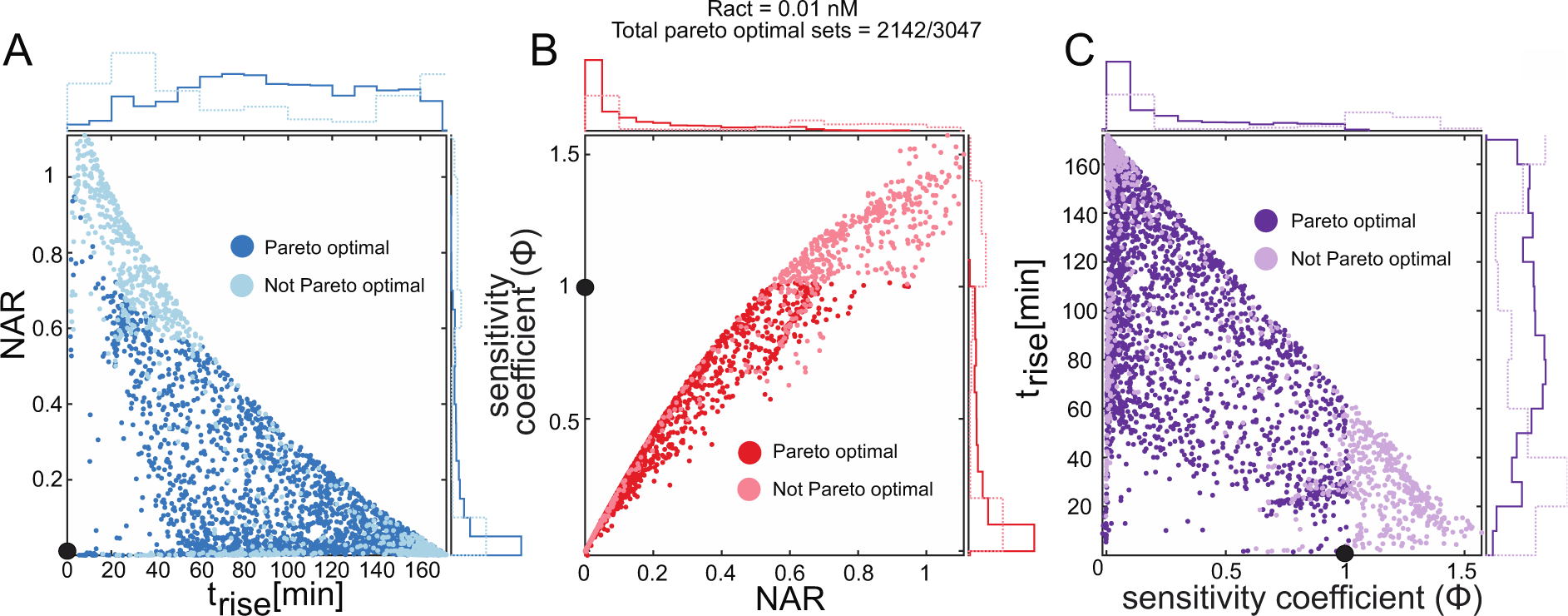
Pareto optimality of POs. (A-C) The projections of the three-dimensional PM-space. Darker filled circles represent Pareto optimal POs, whereas lighter circles represent POs that are not on the Pareto front. The histogram with solid lines indicates the distribution of Pareto optimal POs and the histogram with dashed line indicates the distribution of POs that are not Pareto optimal. **(A)** The t_rise_-NAR projection of the three-dimensional manifold. **(B)** The NAR-sensitivity coefficient (φ) projection of the three-dimensional manifold. **(C)** The sensitivity coefficient (φ) - t_rise_ projection of the three-dimensional. The optimal point (UP) is indicated with black filled circles.

### Mapping performance of biological systems to NCPs

To correlate different system behaviors with values of NCPs, we considered three different potential system behaviors. In the first behavior, System 1 favors linearity and rise time (emphasizing POs 1 and 3) while sacrificing NAR (Fig. 7A). In the second behavior, System 2 favors a small NAR (emphasizing PO2), while sacrificing rise time and linearity (Fig. 7A). In the final behavior, System 3 favors a short response time and high noise filtering (emphasizing POs 1 and 2) while sacrificing linearity (Fig. 7A). In these three different regions of PM-space, being limited to the Pareto Front, the NCPs take on hallmark distributions (Fig. 7B-P). For example, System 1 is characterized by high phosphatase and Smad1 levels, and low Smad4 levels. System 2 has low phosphatase and medium Smad1 and Smad4 levels. Finally, System 3 has medium phosphatase and Smad4 levels, and high Smad1 levels. This relationship between NCPs and PMs would allow us to predict NCP values from experimentally-observed systems-level behavior and, perhaps more importantly, to rationally control systems-level behavior by manipulating NCP levels.

**Figure 7:**
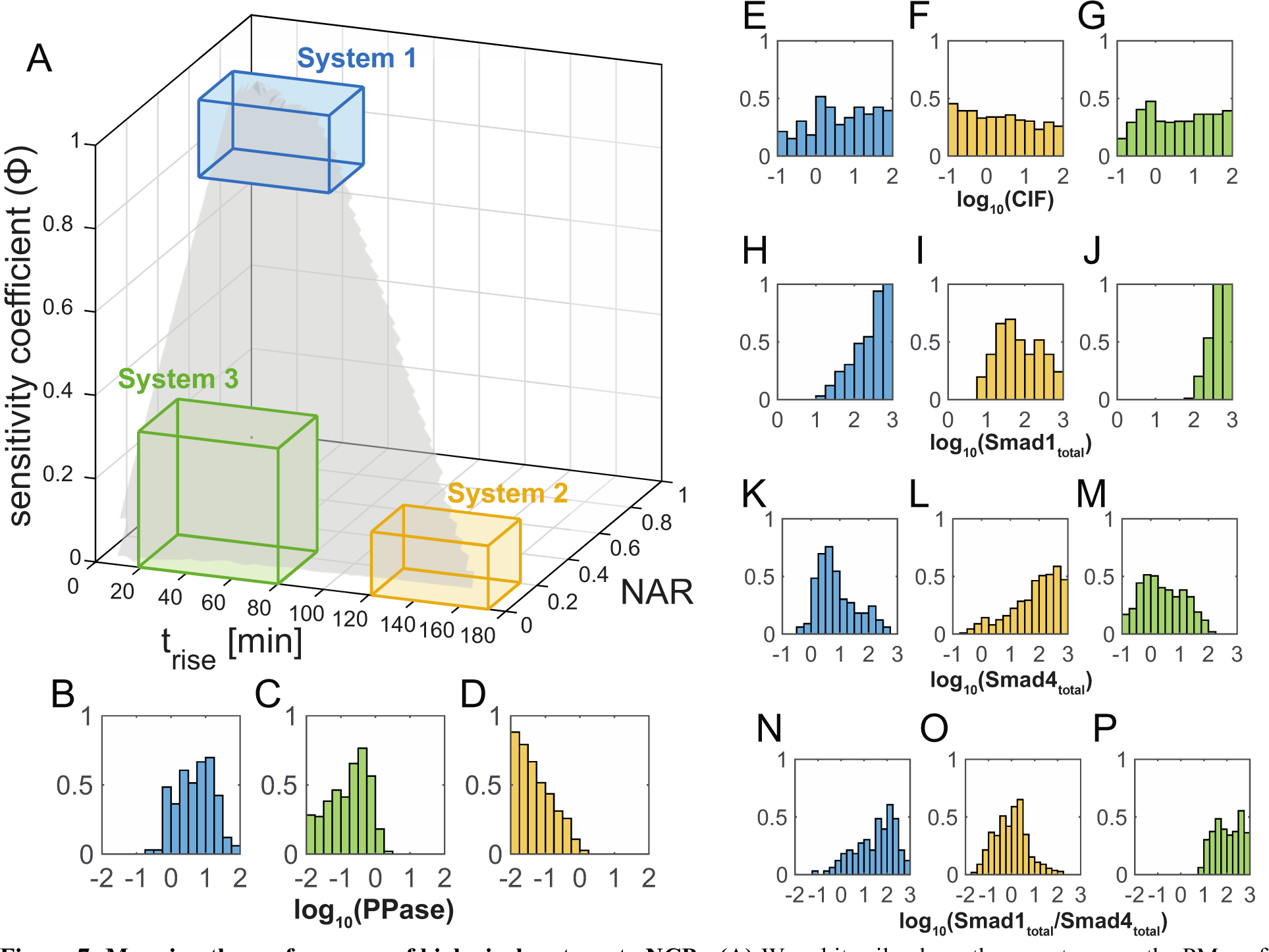
Mapping the performance of biological systems to NCPs. (A) We arbitrarily chose three systems on the PM-surface such that system 1 favors linearity, system 2 favors NAR, and system 3 favors both rise time and NAR. The three systems lie on different regions on the surface fit through all the Pareto optimal PMs. **(B-D)** The distribution of PPase, normalized by a probability distribution function, for the three systems in (A). **(B)** The behavior in system 1 is possible due to high PPase values. **(C)** The behavior in system 2 is possible due to intermediate PPase values. **(D)** The behavior in system 3 is possible due to low PPase values. **(E-G)** The distribution of CIF, normalized by a probability distribution function, for the three systems in (A). **(E)** CIF has a uniform distribution over the entire range. **(F)** CIF has a uniform distribution over the entire range. **(G)** CIF has a uniform distribution over the entire range. **(H-J)** The distribution of Smad1_total_, normalized by a probability distribution function, for the three systems in (A). **(H)** The behavior in system 1 is possible due to high Smad1_total_ values. **(I)** The behavior in system 2 is possible due to intermediate Smad1_total_ values. **(J)** The behavior in system 3 is possible due to very high Smad1_total_ values. **(K-M)** The distribution of Smad4_total_, normalized by a probability distribution function, for the three systems in (A). **(K)** The behavior in system 1 is possible due to low Smad4_total_values. **(L)** The behavior in system 2 is possible due to high Smad4_total_ values. **(M)** The behavior in system 3 is possible due to intermediate Smad4_total_ values. **(N-P)** The distribution of the Smad1_total_/Smad4_total_, normalized by a probability distribution function, for the three systems in (A). **(N)** The behavior in system 1 is possible due to high Smad1_total_/Smad4_total_ values. **(O)** The behavior in system 2 is possible due to low Smad1_total_/Smad4_total_ values. **(P)** The behavior in system 3 is possible due to very high Smad1_total_/Smad4_total_ values.

### Phosphatase levels are variable across species

Our results have shown that nuclear phosphatase, which dephosphorylates PSmad, has the greatest effect of the NCPs on the balance of trade-offs among the POs. To ascertain the variability in phosphatase levels across different species, we used available RNA-Seq data from *Drosophila* embryos, zebrafish embryos, and human cell data. Several phosphatases have been previously linked to the Smad pathway, such as PPM1A [33–35], PPM1H [36], MTMR4 [37], SCP1/2/3[38], SCP4/CTDSPL2 [39], Dullard, Pyruvate dehydrogenase phosphatase (PDP) [40], PP2A [41], etc. Notably, we identified that PPM1A, MTMR4, and Dullard were consistently expressed in *Drosophila*, zebrafish, and human cells. We focused on the available published RNA-Seq data for zebrafish embryos [42], *Drosophila* embryos [43], and human aorta cells [44] specifically examining the expression of the phosphatase Dullard (ctdnep1a in zebrafish and ctdnep1 in humans). The current data reflected that Dullard mRNA exhibited significant expression during the developmental stages of both zebrafish and *Drosophila* embryos, where the BMP/Smad pathway plays a crucial role in shaping the major body axis for these two species. The RNA-Seq dataset is limited in regulated by individual experiment processes, making direct comparisons across species on a universal scale challenging [45]. For a broader perspective on expression levels throughout different species, we conducted a comparison of Dullard expression relative to two highly conserved housekeeping genes, GAPDH (glyceraldehyde-3-phosphate dehydrogenase) and UBC (Ubiquitin C) (Data summary available in Supplementary Text Section 5). Our findings indicate that the relative expression level of Dullard is higher in *Drosophila* embryos than in zebrafish embryos as measured against both GAPHD and UBC. Our findings are consistent with our prior modeling outcomes, and a higher level of PPase maintains POs by achieving a balance between lower noise filtration and heightened sensitivity, leading to quicker response times.

## Discussion

Highly conserved cell-cell signaling pathways, such as the BMP pathway, operate within multiple biological systems, and as such, must be adaptable and versatile to achieve a variety of goals specific to each organism. A fundamental question is how the general morphogen pathway has adapted to the specific needs of a given system, and what mechanisms are at its core in fine-tuning system performance. In the case of BMP-Smad pathway, despite the central and highly conserved role that BMP signaling plays in tissue patterning, the dynamics of signal transmission from the BMP input to the PSmad output varies widely across taxa or developmental stage. For example, BMP signaling requires a rapid response in *Drosophila* and zebrafish embryo development, which occur on the order of minutes to a few hours. On the other hand, in *Drosophila* pupal vein formation and hiPSC differentiation, BMP signaling persists for hours and days, implying that noise filtering may be more important than a rapid response [22]. These observations suggest that the shared BMP signaling module may be tuned to emphasize different POs depending on the context. In certain scenarios, an “ideal” system with optimal POs would respond rapidly, filter noise, and act as a linear sensor of BMP concentration. These response features play a crucial role in determining the fitness of an organism when it faces challenges such as biological noise and perturbations. However, real communication systems are subject to constraints and must navigate trade-offs between various optimal solutions. Balancing these competing factors becomes essential for the functional adaptation and robustness of the BMP-Smad pathway.

Here we investigated the effect of non-conserved parameters (NCPs) on the performance objectives (POs) of the Smad nucleo-cytoplasmic dynamic model. The investigation involved the evaluation of three specific PMs: response time (PM1), noise-amplification-ratio (NAR) (PM2), and sensitivity coefficient (PM3). To assess these objectives, we evaluated the POs by characterizing the response of the system to the localization of (pSmad1)_2_/Smad4 in the nucleus upon pathway activation. The results revealed an intriguing relationship between the concentration of one NCP, PPase, and the POs. As the concentration of PPase increased, the response time of the system decreased. This suggests that a higher concentration of PPase led to a more rapid response, indicating its role in modulating the response timing of the Smad nucleo- cytoplasmic dynamics. On the other hand, the results also unveiled that an elevated concentration of PPase correlated with an increased noise-amplification capability. This implies that PPase plays a vital role in filtering out extraneous noise, enhancing the system’s ability to maintain accurate signaling despite external perturbations. Additionally, as the concentration of PPase was increased, the sensitivity coefficient of the system also increased. This suggests that PPase concentration influences the system’s sensitivity to variations in the input signals, potentially allowing for more precise and nuanced responses to changes in the cellular environment. PPase plays an essential role in regulating the TGF-β/BMP signaling pathway by dephosphorylating key components to modulate its activity. Based on our preliminary analysis of the available published RNA-Seq data from *Drosophila* and zebrafish embryos, as well as human cell data, the phosphatase expression levels vary across different species, the relative expression level of Dullard is higher in *Drosophila* embryos than in zebrafish embryos as measured against both GAPHD and UBC. Our findings are consistent with our prior modeling outcomes, and a higher level of PPase maintains POs by achieving a balance between lower noise filtration and heightened sensitivity, leading to quicker response times. These observations align with the developmental requirements of *Drosophila* embryos, which necessitate rapid responsiveness during their developmental stages. The findings underscore the significance of protein phosphatases in modulating TGF-β/BMP signaling and highlight the need for further exploration into their role.

While PPase levels play the largest role in shaping the balance of trade-offs among the POs we analyzed here, it is only a singular parameter, variation of PPase alone can, at best, trace out a single curve in PO- space. Therefore, the other NCPs, in particular, Smad1 and Smad4 concentrations, can fine-tune the balance of trade-offs to allow the system to sample the entire Pareto surface. Interestingly, our model predicts that Smad1 and Smad4 have largely opposite effects on the POs. Our model predictions of the relationship between NCPs and POs could be tested experimentally in a variety of BMP systems across multiple animal species.

In this work, we focused on only three POs, given they are requirements for communications systems [23, 24, 31, 46]. However, the BMP system may have other competing POs that must also be balanced depending on the context. For example, signal amplification, precision, and information processing are plausible POs that could also be considered in the future [23, 24, 47].

This study presents a fresh perspective on the regulation of BMP/Smad signaling by emphasizing the significance of considering concentration-dependent effects in pathway regulation across different species. This research also sheds light on the intricate mechanisms that underlie the adaptability and robustness of the BMP-Smad pathway, offering valuable insights into its regulation and plasticity. Further exploration is warranted to uncover the mechanisms governing the concentrations of pathway components and to understand their implications in the fields of developmental biology, disease-related contexts, and regenerative medicine.

## Methods

### Simulating the Smad model

For this study, we modified the Smad signal transduction model reported in Schmierer et al., 2008 [27]. In this model, the ligand TGF-β binds to the receptor resulting in the phosphorylation of Smad2, a receptor-regulated Smad (R-Smad). The phosphorylated-Smad2 (pSmad2) forms homomeric or heteromeric complexes with Smad4, a common mediator Smad (Co-Smad) which then translocates to the nucleus to regulate gene expression. We implemented this mathematical model to simulate the BMP pathway and model Smad1 dynamics as its network structure is similar to TGF- β pathway.

It has been reported in the literature [48–50] that the active signaling complex is a trimer consisting of two molecules of pSmad1 and one molecule of Smad4. We accordingly modified (see Supplementary Text Section 3) the model to account for interactions between dimers that result in the formation of the trimer — (pSmad1)_2_/Smad4. We fit this modified model to the nuclear EGFP-Smad1 concentration data reported in [27].

In our modified model (referred to as the Smad model), before the activation of the pathway, only monomeric-Smad1 is present in the nucleus, and after the pathway is activated due to TGF-β signaling all Smad species such as pSmad1, pSmad1/Smad4, pSmad1/pSmad1 and (pSmad1)_2_/Smad4 are present in the nucleus. This leads to the accumulation of Smad1 species in the nucleus when the TGF-β signaling is active. All the Smad species (monomeric and dimeric) can dynamically move between the cytoplasm and the nucleus. An externally supplied inhibitor that deactivates the receptors turns off this signaling network.

### Simulation conditions

The initial conditions of all protein complexes were set to 0.

To calculate the POs at randomly generated NCPs we simulated the Smad model for 24h (R_max_) and 48h (R_min_).

Smad1 and Smad4 initial conditions

Smad1 distribution in initial conditions:

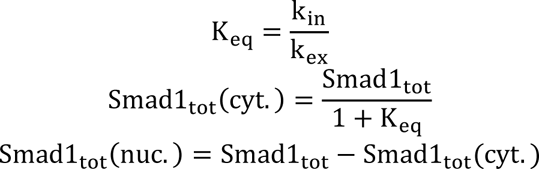

Smad4 is equally distributed in the cytoplasm and the nucleus before the activation of the pathway.

CIF and PPase were passed to the model without modification.

### Parameters for Smad model

We implemented Improved Stochastic Ranking Evolutionary Strategy plus (ISRES+) [51] to estimate the 10 variable parameters in the model. The model was fit to nuclear EGFP-Smad1 concentrations in HaCat cells in the presence of TGF-β signaling and later when an inhibitor of the Smad pathway was externally added. ISRES+ is a (μ-λ)-based evolutionary algorithm employed to solve global optimization problems using stochastic ranking. It additionally has gradient-based algorithms embedded to improve the search strategy, called lin-step and newton-step. The algorithm was run for 100 generations with 150 individuals, with a recombination rate of 0.85. Lin-step contributed one individual and newton-step contributed two individuals to every generation. The complete description of hyperparameters supplied to ISRES+ is in the Supplementary Text Section 3. The best individual of all generations, i.e., the parameter set with the best fitness score was chosen for all the simulations in this study. Additional details are in the Supplementary Text Section 3.

### Receptor concentrations

The activated receptor level inputs used to calculate the PM-2 were obtained using a model from Larson et al., [30]. This model uses the GillesPy python package [52] to simulate the cell surface BMP receptor network as it interacts with given extracellular BMP ligand concentrations. There were two sets of starting parameters for the number of receptor components. Based on literature TGF-β receptor estimations from [53, 54], we used a value of 14,000 TGF-β receptors per cell split between type-I and type-II. [55] suggests that at some stages only ten percent of these may be present on the cell membrane. To this end, the simulations were run a second time with “R_min_” being ten percent of “R_max_”. For each of the two starting receptor levels, extracellular BMP ligand concentrations ranging from 0.001 to 3 nM were used to initialize the simulation. Following 24 to 48 hours of simulation, all species population values were tracked over 24 hours to collect noise information at the steady state.

The R_act_ values for “R_max_” range from 0.01 nM to 3.5 nM, and for “R_min_” range from 0.001 to 0.1 nM (Fig S5).

To calculate the PM-1 and PM-3, we adapt (details in the Supplementary Text Section 4) our model and simulate at starting receptor levels as a constant input.

### Sampling non-conserved protein concentrations

We generated 10^4^ sets of three non-conserved protein concentrations (Smad4, CIF and PPase) using Latin-hypercube design with a sample density of 10^4^. We uniformly sample the Smad1/Smad4 ratio in the logspace.

The following were the range we generated to sample within:

|  | Min (nM) | Max (nM) |
| --- | --- | --- |
| CIF | $10^{-1}$ | $10^2$ |
| PPase | $10^{-2}$ | $10^2$ |
| Smad1<sub>tot</sub> | $10^{-1}$ | $10^3$ |
| Smad4<sub>tot</sub> | $10^{-1}$ | $10^3$ |

The simulation results with nuclear (pSmad1)_2_/Smad4 concentration less than 0.1 nM are not shown in the figures since these concentrations are too low to be biologically relevant.

### Calculating the three Performance Objectives

The three performance objectives (POs) are:

#### 1. Rise time (t_rise_)

The rise time is defined as the duration of a response to reach 95% of its steady state value. We calculated the rise time using a MATLAB function stepinfo from the control systems toolbox, on the dynamic profile of total nuclear pSmad14. The steady state pS2S4 concentration was externally supplied as an input to stepinfo, which was calculated by solving the system of ODEs using newton’s method with the final condition from ode23tb in MATLAB as an initial guess. The ‘RiseTimeLimits’ were defined as [0 0.95]

#### 2. Noise amplification ratio (NAR)

The NAR is defined as the ratio of coefficient of variation of the response signal to the input signal. We used the steady state of nuclear (pSmad1)_2_/Smad4 obtained in PO1 as an initial condition to simulate continuous model again with a stochastic receptor input. We simulate the model for the same timespan as PO1. The NAR is then calculated as the ratio CV of (pSmad1)_2_/Smad4 over R_act_:

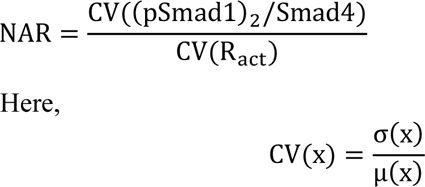

We calculated the sensitivity coefficient at the steady state concentration of (pSmad1)_2_/Smad4 obtained by solving the system of ODEs using newton’s method with the final condition from ode23tb in MATLAB as an initial guess. The sensitivity coefficient defined as:

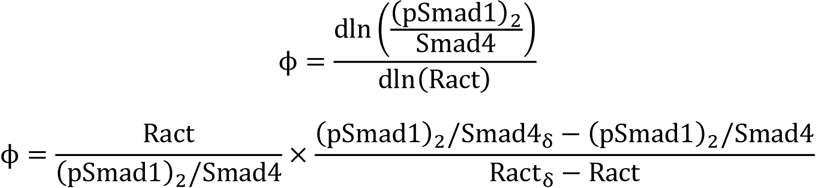

Here, Ract_δ_ = 1.01R_act_ and (pSmad1)_2_/Smad4_δ_ is the steady state concentration of (pSmad1)_2_/Smad4 for Ract_δ_.

### Pareto optimality

We define an optimal point that has coordinates such that t_rise_ = 0, NAR = 0 and ϕ = 1.

We determined pareto optimality by selecting non-dominated solutions by calculating the distance between every point from every other point on the PM surface. These distances were calculated by normalizing the POs to ensure they lie in the same range. The response speed, t_rise_ (PM 1) was normalized by the maximum t_rise_ in the random screen. The abs(abs(ϕ) ) value of φ was used to ensure φ = 1 is optimal.

### Centroid of point clouds

The POs were sorted into ten discrete clusters to visualize the effect of varying NCP. We calculated the centroid of each cluster by taking the mean of the three POs. The same procedure is used to calculate the centroid of point cloud clusters at all R_act_ concentrations.

The Euclidean distance (d) was calculated as follows:

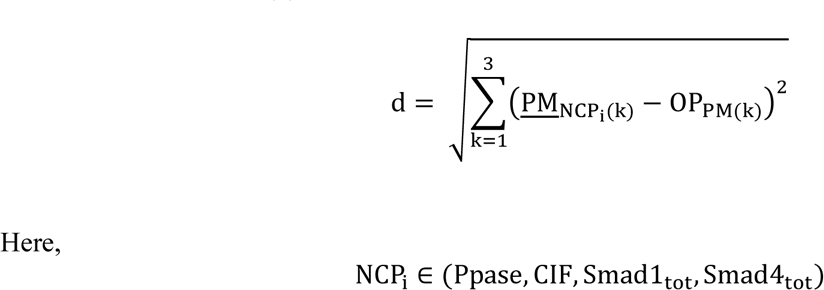

### Surface Fitting and generating distribution of NCPs

To visualize the PM-space in three dimensions and correlate system behavior using a data-driven approach, we fit a curve through all the points on the Pareto using the “fit” function in MATLAB using ‘cubicinterp’ method and setting ‘ExtrapolationMethod’ to ‘none’. The resulting surface is as shown in Fig. 7A. To generate the distribution of NCPs in distinct regions on the PM-surface, we draw a cuboid in three dimensions using the interactive MATLAB function ‘drawcuboid’, and reverse calculate the distribution of NCPs that allow each system behavior. The histograms were normalized by probability distribution function.

## Acknowledgments

This work is based upon efforts supported by the EMBRIO Institute, contract #2120200, a National Science Foundation (NSF) Biology Integration Institute.

This research was supported in part by the NIH grants R01GM132501 awarded to D.U. and by NSF grant 2313692 awarded to G.T.R. and by NSF grant 2047903 awarded to D.H.-P.

Portions of this research were conducted with the advanced computing resources provided by Texas A&M High Performance Research Computing.

## References

1. Shi Y, Massagué J. Mechanisms of TGF-β signaling from cell membrane to the nucleus. cell. 2003;113(6):685–700.

2. Bayat V, Jaiswal M, Bellen HJ. The BMP signaling pathway at the Drosophila neuromuscular junction and its links to neurodegenerative diseases. Current Opinion in Neurobiology. 2011;21(1):182–8. doi: 10.1016/j.conb.2010.08.014.

3. Davies EL, Fuller MT. Regulation of Self-renewal and Differentiation in Adult Stem Cell Lineages: Lessons from the Drosophila Male Germ Line. Cold Spring Harbor Symposia on Quantitative Biology. 2008;73(0):137–45. doi: 10.1101/sqb.2008.73.063.

4. Shaikh S, Ravenndranath R, Banerjee M, Joseph A, Jahgirdar P. Evidence for transforming Growth Factor-Beta 3 gene polymorphism in non-syndromic cleft lip and palate patients from Indian sub-continent. Medicina Oral Patología Oral y Cirugia Bucal. 2012:e197–e200. doi: 10.4317/medoral.17453.

5. Finnson KW, Chi Y, Bou-Gharios G, Leask A, Philip A. TGF-b signaling in cartilage homeostasis and osteoarthritis. Front Biosci (Schol Ed). 2012;4(2):251e68.

6. Samanta D, Datta PK. Alterations in the Smad pathway in human cancers. Frontiers in bioscience (Landmark edition). 2012;17:1281.

7. Restrepo S, Zartman J, Jeremiah, Basler K. Coordination of Patterning and Growth by the Morphogen DPP. Current Biology. 2014;24(6):R245–R55. doi: 10.1016/j.cub.2014.01.055.

8. Bier E, De Robertis EM. BMP gradients: A paradigm for morphogen-mediated developmental patterning. Science. 2015;348(6242):aaa5838.

9. Raftery LA, Twombly V, Wharton K, Gelbart WM. Genetic screens to identify elements of the decapentaplegic signaling pathway in Drosophila. Genetics. 1995;139(1):241–54.

10. Padgett RW, St. Johnston RD, Gelbart WM. A transcript from a Drosophila pattern gene predicts a protein homologous to the transforming growth factor-β family. Nature. 1987;325(6099):81-4. doi: 10.1038/325081a0.

11. Arora K, Levine MS, O’Connor MB. The screw gene encodes a ubiquitously expressed member of the TGF-beta family required for specification of dorsal cell fates in the Drosophila embryo. Genes & Development. 1994;8(21):2588–601. doi: 10.1101/gad.8.21.2588.

12. Raftery LA, Sutherland DJ. Gradients and thresholds: BMP response gradients unveiled in Drosophila embryos. TRENDS in Genetics. 2003;19(12):701–8.

13. O’Connor MB, Umulis D, Othmer HG, Blair SS. Shaping BMP morphogen gradients in the *Drosophila*embryo and pupal wing. Development. 2006;133(2):183-93. doi: 10.1242/dev.02214.

14. Gavin-Smyth J, Wang Y-C, Butler I, Ferguson L, Edwin. A Genetic Network Conferring Canalization to a Bistable Patterning System in Drosophila. Current Biology. 2013;23(22):2296–302. doi: 10.1016/j.cub.2013.09.055.

15. Asafen HYA, Beseli A, Hiremath S, Williams CM, Reeves GT. Dynamics of BMP signaling in the early *Drosophila* embryo. bioRxiv. 2022:2022.10.20.513072. doi: 10.1101/2022.10.20.513072.

16. Yan SJ, Zartman JJ, Zhang M, Scott A, Shvartsman SY, Li WX. Bistability coordinates activation of the EGFR and DPP pathways in *Drosophila* vein differentiation. Molecular Systems Biology. 2009;5(1):278. doi: 10.1038/msb.2009.35.

17. Tucker JA, Mintzer KA, Mullins MC. The BMP Signaling Gradient Patterns Dorsoventral Tissues in a Temporally Progressive Manner along the Anteroposterior Axis. Developmental Cell. 2008;14(1):108–19. doi: 10.1016/j.devcel.2007.11.004.

18. Ramel M-C, Hill CS. The ventral to dorsal BMP activity gradient in the early zebrafish embryo is determined by graded expression of BMP ligands. Developmental biology. 2013;378(2):170–82.

19. Subileau M, Merdzhanova G, Ciais D, Collin-Faure V, Feige J-J, Bailly S, et al. Bone Morphogenetic Protein 9 Regulates Early Lymphatic-Specified Endothelial Cell Expansion during Mouse Embryonic Stem Cell Differentiation. Stem Cell Reports. 2019;12(1):98–111. doi: 10.1016/j.stemcr.2018.11.024.

20. Derynck R, Akhurst RJ. BMP-9 balances endothelial cell fate. Proceedings of the National Academy of Sciences. 2013;110(47):18746–7. doi: 10.1073/pnas.1318346110.

21. Ponomarev LC, Ksiazkiewicz J, Staring MW, Luttun A, Zwijsen A. The BMP Pathway in Blood Vessel and Lymphatic Vessel Biology. International Journal of Molecular Sciences. 2021;22(12):6364. doi: 10.3390/ijms22126364.

22. Alderfer L, Wei A, Hanjaya-Putra D. Lymphatic Tissue Engineering and Regeneration. Journal of Biological Engineering. 2018;12(1). doi: 10.1186/s13036-018-0122-7.

23. Lander AD, Lo W-C, Nie Q, Wan FYM. The Measure of Success: Constraints, Objectives, and Tradeoffs in Morphogen-mediated Patterning. Cold Spring Harbor Perspectives in Biology. 2009;1(1):a002022-a. doi: 10.1101/cshperspect.a002022.

24. Lo W-C, Zhou S, Wan FY-M, Lander AD, Nie Q. Robust and precise morphogen-mediated patterning: trade-offs, constraints and mechanisms. Journal of The Royal Society Interface. 2015;12(102):20141041. doi: 10.1098/rsif.2014.1041.

25. Qiao L, Zhao W, Tang C, Nie Q, Zhang L. Network Topologies That Can Achieve Dual Function of Adaptation and Noise Attenuation. Cell Systems. 2019;9(3):271–85.e7. doi: 10.1016/j.cels.2019.08.006.

26. Hornung G, Barkai N. Noise Propagation and Signaling Sensitivity in Biological Networks: A Role for Positive Feedback. PLoS Computational Biology. 2008;4(1):e8. doi: 10.1371/journal.pcbi.0040008.

27. Schmierer B, Tournier AL, Bates PA, Hill CS. Mathematical modeling identifies Smad nucleocytoplasmic shuttling as a dynamic signal-interpreting system. Proceedings of the National Academy of Sciences of the United States of America. 2008;105(18):6608–13. doi: 10.1073/pnas.0710134105. PubMed PMID: WOS:000255841600019.

28. Shoval O, Sheftel H, Shinar G, Hart Y, Ramote O, Mayo A, et al. Evolutionary Trade-Offs, Pareto Optimality, and the Geometry of Phenotype Space. Science. 2012;336(6085):1157-60. doi: doi:10.1126/science.1217405.

29. Schuetz R, Zamboni N, Zampieri M, Heinemann M, Sauer U. Multidimensional Optimality of Microbial Metabolism. Science. 2012;336(6081):601-4. doi: doi:10.1126/science.1216882.

30. Larson NJ, Madamanchi A, Karim MS, Li L, Umulis DM. Stochastic Analysis of BMP Receptor Oligomerization During Embryogenesis. [Unpublished Manuscript].

31. Karim MS, Buzzard GT, Umulis DM. Secreted, receptor-associated bone morphogenetic protein regulators reduce stochastic noise intrinsic to many extracellular morphogen distributions. J R Soc Interface. 2012;9(70):1073–83. Epub 20111019. doi: 10.1098/rsif.2011.0547. PubMed PMID: 22012974; PubMed Central PMCID: PMCPMC3306646.

32. Boada Y, Reynoso-Meza G, Picó J, Vignoni A. Multi-objective optimization framework to obtain model-based guidelines for tuning biological synthetic devices: an adaptive network case. BMC Systems Biology. 2016;10(1). doi: 10.1186/s12918-016-0269-0.

33. Lin X, Duan X, Liang Y-Y, Su Y, Wrighton KH, Long J, et al. PPM1A Functions as a Smad Phosphatase to Terminate TGFβ Signaling. Cell. 2006;125(5):915–28. doi: 10.1016/j.cell.2006.03.044.

34. Duan X, Liang Y-Y, Feng X-H, Lin X. Protein Serine/Threonine Phosphatase PPM1A Dephosphorylates Smad1 in the Bone Morphogenetic Protein Signaling Pathway. Journal of Biological Chemistry. 2006;281(48):36526–32. doi: 10.1074/jbc.m605169200.

35. Kokabu S, Nojima J, Kanomata K, Ohte S, Yoda T, Fukuda T, et al. Protein phosphatase magnesium-dependent 1A–mediated inhibition of BMP signaling is independent of Smad dephosphorylation. Journal of Bone and Mineral Research. 2010;25(3):653–60. doi: 10.1359/jbmr.090736.

36. Shen T, Sun C, Zhang Z, Xu N, Duan X, Feng X-H, et al. Specific control of BMP signaling and mesenchymal differentiation by cytoplasmic phosphatase PPM1H. Cell Research. 2014;24(6):727–41. doi: 10.1038/cr.2014.48.

37. Yu J, He X, Chen Y-G, Hao Y, Yang S, Wang L, et al. Myotubularin-related Protein 4 (MTMR4) Attenuates BMP/Dpp Signaling by Dephosphorylation of Smad Proteins. Journal of Biological Chemistry. 2013;288(1):79–88. doi: 10.1074/jbc.m112.413856.

38. Knockaert M, Sapkota G, Alarcón C, Massagué J, Brivanlou AH. Unique players in the BMP pathway: Small C-terminal domain phosphatases dephosphorylate Smad1 to attenuate BMP signaling. Proceedings of the National Academy of Sciences. 2006;103(32):11940–5. doi: 10.1073/pnas.0605133103.

39. Zhao Y, Xiao M, Sun B, Zhang Z, Shen T, Duan X, et al. C-terminal Domain (CTD) Small Phosphatase-like 2 Modulates the Canonical Bone Morphogenetic Protein (BMP) Signaling and Mesenchymal Differentiation via Smad Dephosphorylation. Journal of Biological Chemistry. 2014;289(38):26441–50. doi: 10.1074/jbc.m114.568964.

40. Chen HB, Shen J, Ip YT, Xu L. Identification of phosphatases for Smad in the BMP/DPP pathway. Genes & Development. 2006;20(6):648–53. doi: 10.1101/gad.1384706.

41. Bengtsson L, Schwappacher R, Roth M, Boergermann JH, Hassel S, Knaus P. PP2A regulates BMP signalling by interacting with BMP receptor complexes and by dephosphorylating both the C-terminus and the linker region of Smad1. Journal of Cell Science. 2009;122(8):1248–57. doi: 10.1242/jcs.039552.

42. White RJ, Collins JE, Sealy IM, Wali N, Dooley CM, Digby Z, et al. A high-resolution mRNA expression time course of embryonic development in zebrafish. Elife. 2017;6. Epub 20171116. doi: 10.7554/eLife.30860. PubMed PMID: 29144233; PubMed Central PMCID: PMCPMC5690287.

43. Graveley BR, Brooks AN, Carlson JW, Duff MO, Landolin JM, Yang L, et al. The developmental transcriptome of Drosophila melanogaster. Nature. 2011;471(7339):473-9. doi: 10.1038/nature09715.

44. Lonsdale J, Thomas J, Salvatore M, Phillips R, Lo E, Shad S, et al. The Genotype-Tissue Expression (GTEx) project. Nature Genetics. 2013;45(6):580–5. doi: 10.1038/ng.2653.

45. Deshpande D, Chhugani K, Chang Y, Karlsberg A, Loeffler C, Zhang J, et al. RNA-seq data science: From raw data to effective interpretation. Frontiers in Genetics. 2023;14. doi: 10.3389/fgene.2023.997383.

46. Reeves GT. The engineering principles of combining a transcriptional incoherent feedforward loop with negative feedback. Journal of Biological Engineering. 2019;13(1). doi: 10.1186/s13036-019-0190-3.

47. Dubuis JO, Tkačik G, Wieschaus EF, Gregor T, Bialek W. Positional information, in bits. Proceedings of the National Academy of Sciences. 2013;110(41):16301–8. doi: 10.1073/pnas.1315642110.

48. Schmierer B, Hill CS. TGF beta-SMAD signal transduction: molecular specificity and functional flexibility. Nature Reviews Molecular Cell Biology. 2007;8(12):970–82. doi: 10.1038/nrm2297. PubMed PMID: WOS:000251173600010.

49. Massagué J, Seoane J, Wotton D. Smad transcription factors. Genes & Development. 2005;19(23):2783-810. doi: 10.1101/gad.1350705.

50. Tiago Gomes and Pau Martin-Malpartida and Lidia Ruiz and Eric Aragón and Tiago NCaMJM. Conformational landscape of multidomain SMAD proteins. Computational and Structural Biotechnology Journal. 2021;19:5210–24. doi: 10.1016/j.csbj.2021.09.009.

51. Bandodkar P, Shaikh R, Reeves GT. ISRES+: an improved evolutionary strategy for function minimization to estimate the free parameters of systems biology models. Bioinformatics. 2023;39(7). doi: 10.1093/bioinformatics/btad403.

52. Abel JH, Drawert B, Hellander A, Petzold LR. GillesPy: A Python Package for Stochastic Model Building and Simulation. IEEE Life Sciences Letters. 2016;2(3):35–8. doi: 10.1109/lls.2017.2652448.

53. Wakefield LM, Smith DM, Masui T, Harris CC, Sporn MB. Distribution and modulation of the cellular receptor for transforming growth factor-beta. The Journal of cell biology. 1987;105(2):965–75. doi: 10.1083/jcb.105.2.965.

54. Nicklas DaSL. Computational modelling of Smad-mediated negative feedback and crosstalk in the TGF-β superfamily network. Journal of The Royal Society Interface. 2013;10(86):20130363. doi: 10.1098/rsif.2013.0363.

55. Chung S-W, Miles FL, Sikes RA, Cooper CR, Farach-Carson MC, Ogunnaike BA. Quantitative Modeling and Analysis of the Transforming Growth Factor β Signaling Pathway. Biophysical Journal. 2009;96(5):1733–50. doi: 10.1016/j.bpj.2008.11.050.

